# Differential effect of anesthesia on visual cortex neurons with diverse population coupling

**DOI:** 10.1101/2020.06.13.150284

**Authors:** Heonsoo Lee, Sean Tanabe, Shiyong Wang, Anthony G. Hudetz

**Affiliations:** Center for Consciousness Science, Department of Anesthesiology, University of Michigan, Ann Arbor, MI, USA, 48105

## Abstract

**Introduction:** Firing rate (FR) and population coupling (PC) are intrinsic properties of cortical neurons. Neurons with different FR and PC have diverse excitability to stimulation, tuning curve, and synaptic plasticity. Therefore, investigation of the effect of anesthesia on neurons with different FR and PC would be important to understand state-dependent information processing in neuronal circuits.

**Methods:** To test how anesthesia affects neurons with diverse PC and FR, we measured single-unit activities in deep layers of primary visual cortex at three levels of anesthesia with desflurane and in wakefulness. Based on PC and FR in wakefulness, neurons were classified into three distinct groups: high PC-high FR (HPHF), low PC-high FR (LPHF), and low PC-low FR (LPLF) neurons.

**Results:** Applying repeated light flashes as visual stimuli, HPHF neurons showed the strongest early response (FR at 20-150ms post-stimulus) among the three groups, whereas the response of LPHF neurons persisted longest (up to 440ms). Anesthesia profoundly altered PC and FR, and differently affected the three neuron groups: (i) PC and FR became strongly correlated suppressing population-independent spike activity; (ii) Pairwise correlation of spikes between neurons could be predicted by a PC-based raster model suggesting uniform neuron-to-neuron coupling; (iii) Contrary to evoked-potential studies under anesthesia, the flash-induced early response of HPHF neurons was attenuated, and their spike timing was split and delayed; (iv) Late response (FR at 200-400ms post-stimulus) was suppressed both in HPHF and LPHF neurons.

**Conclusions:** Anesthetic-induced association between PC and FR suggests reduced information content in the neural circuit. Altered response of HPHF neurons to visual stimuli suggests that anesthesia interferes with conscious sensory processing in primary sensory cortex.

## 1. Introduction

Firing rate (FR) and population coupling (PC) are intrinsic properties of cortical neurons. They are thought to be invariant to behavioral conditions such as waking and sleep states, and also to resting and stimulation conditions [1,2]. These properties are also closely related to the information processing functions of each neurons [3,4]. For example, spikes of strongly coupled neurons (choristers) is thought to reflect behavioral state of the animal and driven by nonspecific excitation such as optogenetic stimulation [1,5]. On the other hand, weakly coupled neurons (soloists) appear to respond selectively and precisely to sensory stimuli [4,6]; e.g., in primary visual cortex, these neurons show a precise tuning curve to a grating patch. In addition, it has been reported that sparsely firing neurons showed greater synaptic plasticity during sleep [6].

Previous studies were performed under different behavioral conditions such as waking [1,5,6], sleep [6,7], or under anesthesia [1,4]. No study to-date has investigated the dose-dependent effect of anesthesia on FR and PC properties in the same animal in a systematic manner. Considering the substantial changes in neural network dynamics [8,9] and the disruption in conscious cognitive functions anesthetics cause, we asked if and how anesthesia may affect neurons depending on their PC and FR and how this may disrupt sensory processing. Such an investigation should provide a more precise understanding of the state-dependence of different neuron groups with distinct PC and FR, their role in sensory information processing, and how anesthesia disturbs consciousness.

To address our questions, we recorded single-unit activities from deep layers of primary visual cortex of rats during graded levels of anesthesia with the common clinically used anesthetic desflurane and in wakefulness. PC, FR, and neuron-to-neuron correlation were estimated from spontaneous activity and the interrelationship between these measures was explored. In addition, neuron populations were divided into subgroups based on their PC and FR values. The response of these neuron groups to visual flash stimulation was investigated as a function of the level of anesthesia.

## 2. Methods

### 2.1 Electrode implantation

The study was approved by the Institutional Animal Care and Use Committee in accordance with the Guide for the Care and Use of Laboratory Animals of the Governing Board of the National Research Council (National Academy Press, Washington, D.C., 2011). Adult male Long-Evans rats (n = 8; 300-350 g weight) were housed in a reverse light-dark cycle room for 5-7 days prior to electrode implantation. Ad libitum access for food and water was provided for the duration of the experiment. A sixty-four contact multi-electrode array (silicon probes; shank length 2 mm, width 28-60 µm, probe thickness 15 µm, shank spacing 200 µm, row separation 100 µm, contact size 413 µm2; custom design a8×8_edge_2mm100_200_413, Neuronexus Technologies, Ann Arbor, MI) was chronically implanted in the primary visual cortex of each rat. Electromyogram was recorded using a pair of insulated wires (PlasticsOne, Inc., Roanoke, VA) positioned bilaterally into the nuchal muscles. A craniotomy of rectangular shape of approximately 2 × 4 mm was prepared, the exposed dura mater was resected, and the electrode array was inserted using a micromanipulator to the final position 1.6 mm below the pial surface. The perimeter was covered with silicone gel (Kwik-Sil, World Precision Instruments, Sarasota, FL). Additional sterilized stainless-steel screws were used to secure the electrode to the cranium. The assembly was embedded with dental cement (Stoelting Co., Wood Dale, IL). Carprofren (5 mg/kg s.c. once daily) was administered for three days after surgery. The animals were observed for 7–10 days after surgery for any infection or other complications.

### 2.2 Experimental design

Extracellular recording was performed in a closed, ventilated anesthesia chamber 1-8 days after surgery. Desflurane was administrated with a stepwise decreasing concentration, 6, 4, 2, and 0%. Notice that considering the purpose of this study to explore the effects of anesthetic, the focus in the analysis was on changes from wakefulness to deep anesthesia levels (from 0 to 6%). In order to achieve equilibrium concentration, there was a 15 minutes of transient period between every concentration levels. Each concentration level comprised of resting state and visual stimulation sessions. During resting state session, spontaneous activity was recorded for 20 minutes per each desflurane level. For one experiment which was performed in the beginning of the study, only 10 minutes of spontaneous activity was recorded per anesthetic concentration. Because every neuronal variable was quantified from 10 second epochs, 10 minute data length per desflurane concentration should not affect the final conclusions and the data was kept for the analysis. During anesthesia, spontaneous spike activity of neurons was occasionally desynchronous while showing low firing rate. This paradoxical desynchronized activity in anesthesia is unexpected and has never been reported. Because this phenomena is beyond the purpose of the study, we excluded the desynchronization periods from the analysis; the paradoxical desynchronization brain state was identified based on agglomerative clustering algorithm, and on average, 40.0%, 6.5%, 10.3%, and 0.3% of data was excluded in 6, 4, 2, and 0% desflurane, respectively. The follow-up study will explore the details of this brain state. During visual simulation session, light flashes of 10ms duration were delivered to the retina by transcranial illumination with randomized intervals (2-4 seconds; 100 trials per concentration level). Flashes of 1ms duration were also delivered in six out of eight experiments due to slightly different experimental protocol; the recording in this period was not analyzed. Anesthetic concentration in the holding chamber was continuously monitored (POET IQ2 monitor; Criticare Systems, Inc., Waukesha, WI). Core body temperature was maintained at 37°C by subfloor heating.

### 2.3 Electrophysiological recording and identification of single units

Extracellular potentials were recorded at 30 kHz sampling rate (SmartBox; Neuronexus Technologies, Ann Arbor, MI). The high frequency components of raw data were used for detecting unit activities (neuronal spikes). First, the sixty-four signals were median-referenced. Data with absolute value greater than 10 SD were removed as artifacts. Noticeable noise episodes were visually inspected and manually excluded from the analysis. One experiment was excluded from the analysis due to severe noise contamination (n = 7). Single unit activity was identified using a template-based spike sorting software, Spiking Circus [10]. On average, 36 ± 14 (mean ± SD) single units (neurons) were obtained. All spontaneous activity properties were calculated for non-overlapping 10 second epochs. For analysis of neural response to visual stimuli, recording period ± 1 was used to calculate firing rate (FR). The neurons were divided into putative excitatory (pE) and inhibitory (pI) units based on the spike waveform and autocorrelogram. Units with short half-amplitude width, short trough-to-peak time, and fast-spiking pattern were manually selected as a pI [11] and the rest of the units were classified into PE.

### 2.4 Population coupling analysis

We used the raster marginal model [1] to create a random raster with equivalent PC and FR as in the original raster. The model was applied to 10 second epochs across the data at resolution of 20ms Heaviside step bins with no overlap. To implement the raster marginal model, we extracted from the original raster (i.e., spike train data with ‘1’ for spike and ‘0’ for non-spike) (i) the histogram of the instantaneous population rate by summing ‘1’s through neurons, (ii) the total firing (linearly related to FR) by summing 1s through time, and (iii) the PC by the dot product between the instantaneous population rate and the time series of each neuron. First we created a random raster meeting criteria of (i). From this random raster, criterion (ii) was met by repeatedly switching ‘1’s of neurons with total firing higher than the original raster, with ‘0’s of neurons with total firing lower than the original raster. Finally, we met criterion (iii) by finding ‘1’ and ‘0’ from a neuron with PC higher than the original raster, and from the same time points, switch respectively ‘0’ and ‘1’ of a neuron with PC lower than the original raster. We repeated this process until the error was lower than the total number of neurons.

### 2.5 Statistical analysis

Statistical analyses were conducted using StatsModels library (www.statsmodels.org) in Python 3.7. All properties of 2, 4, and 6% desflurane were compared with those of 0% desflurane (waking state). We used linear mixed models (LMM) based on restricted maximum likelihood estimation. For all LMMs, the desflurane concentration (categorical independent variable) were used as a fixed effect. The random effect included the different animals and individual neurons. Post-hoc pairwise comparisons were made between the four different levels or between three different neuronal groups using a Bonferroni adjusted p-value (in all figures, *: p < 0.05, **: p < 0.01, and ***: p < 0.001).

## 3. Results

### 3.1 PC and FR are increasingly associated in anesthesia

How anesthetic modulates PC and FR of spontaneous activity of individual neurons was examined. First, we classified neurons based on PC and FR values in 0% desflurane (i.e., wakefulness) using K-means clustering. This was different from previous studies in which neurons were classified by either PC or FR alone [4,6–8]. Figure 1A illustrates the clustering into three groups of neurons; high PC-high FR (HPHF; n = 55), low PC-high FR (LPHF; n = 99), and low PC-low FR (LPLF; n = 97), which represent active choristers, active soloists, and inactive soloists, respectively. The clusters began to slightly overlap with an increase of the anesthetic but still remained discernible.

**Figure 1.**
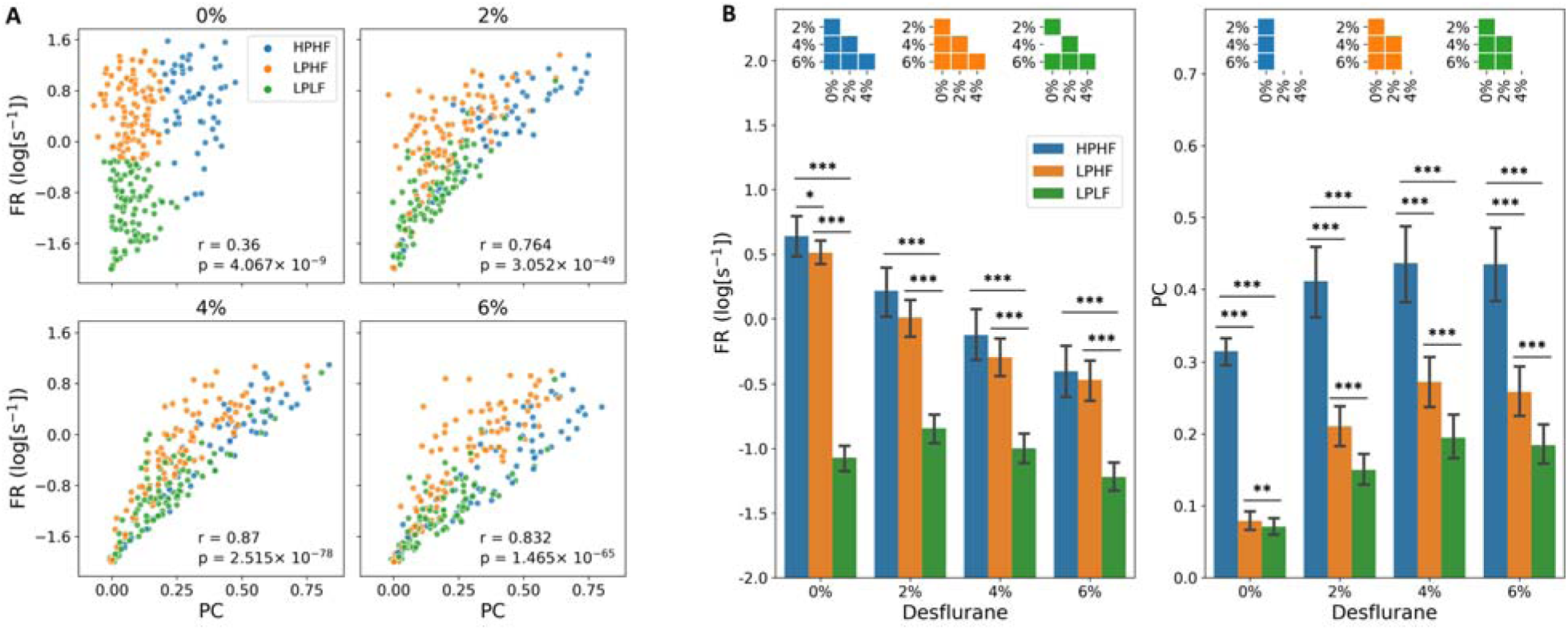
Effect of anesthesia on population coupling (PC) and firing rate (FR) of three classified groups of neurons during spontaneous activity. (A) Cluster plot neurons in three groups classified by K-means clustering with FR and PC as features. Four panels show clusters in wakefulness (0%) and anesthesia (2%, 4%, and 6%) where percent refers to inhaled concentration of desflurane. K-means clustering was performed based at 0% and group identity was kept the same for the other three conditions. Three neuron groups are labeled as high PC-high FR (HPHF, blue), low PC-high FR (LPHF, orange), and low PC-low FR (LPLF, green). Note that PC and FR became more positively correlated in anesthesia; see r (correlation) and corresponding p-value in each panel. (B) Summary of mean FR (left) and PC (right) in wakefulness and anesthesia. Error bars indicate CI 95% from all recorded neurons (n=251). Insets indicate pairwise statistically significant difference among conditions (p < 0.05). Pairwise comparison among neuron groups at each level of anesthesia are indicated by asterisks.

A positive correlation between PC and FR was evident in all four conditions, but it was consistently higher under anesthesia. The cluster of LPHF neurons in particular began to spread in anesthesia, enhancing the correlation of PC and FR. The explained variance by the first principal component of LPHF neurons was 0.216, 0.512, 0.642, and 0.753, for 0, 2, 4, and 6% desflurane, respectively.

Neurons were also classified into putative excitatory (pE) and putative inibitory (pI) types based on their waveform features and autocorrelogram as described in the Methods. The pE neurons comprised 58.2, 91.9, and 95.9% of HPHF, LPHF, and LPLF neurons, respectively. The pI neurons comprised 41.8, 0.81, and 0.41% of HPHF, LPHF, and LPLF neurons, respectively. Thus, in wakefulness, essentially all soloists were pE neurons, whereas choristers were split between pE and pI neurons.

### 3.2 Neurons are constrained to the population

Next, we examined state-dependent changes of PC and FR in each neuron group. FR of HPHF and LPHF neurons gradually decreased as the anesthesia deepened (Fig. 1B left). LPLF neurons showed a similar trend but dropped their FR upon waking. PC showed the opposite trend to FR, it increased in all the three groups (Fig. 1B right). The increase was more pronounced in LPHF and LPLF neurons. Despite the significant changes in both PC and FR, the group identity of neurons was preserved across anesthesia. That is, the PC and FR ranks of the three neuronal groups were the same in all conditions. For individual neurons, Spearman correlation of FR between different levels was, 0.668, 0.566, 0.548, 0.899, 0.860, and 0.906, for 0 vs.2%, 0 vs. 4%, 0 vs. 6%, 2 vs. 4%, 2 vs. 6%, and 4 vs. 6%, respectively. Spearman correlation of PC was, 0.639, 0.509, 0.505, 0.858, 0.776, and 0.882, for 0 vs.2%, 0 vs. 4%, 0 vs. 6%, 2 vs. 4%, 2 vs. 6%, and 4 vs. 6%, respectively. All correlation values were statistically significant (p < 0.05).

### 3.3 Population coupling predicts pairwise correlation

Pairwise correlation of neurons reflects the functional organization of the underlying network. The overall strength of pairwise correlation was increased in anesthesia (Fig 2A, first row). In wakefulness, the correlation among HPHF neurons was stronger than that of the other two groups but such a cluster-like formation diminished as anesthesia deepened. We hypothesized that anesthesia may suppress neuron-to-neuron communication in spite of the increase in synchrony, such that pairwise correlation becomes less complex. If so, FR and PC alone would be able to predict the observed pairwise correlation. To test the hypothesis, we used a spike raster model to predict the pairwise correlation matrix from PC and FR of each neuron and the distribution of the population rate as described in Methods. The predicted correlation matrix visually resembled the observed correlation matrix (Fig. 2A, second row). In order to estimate the agreement between predicted and observed correlation, the normalized difference was calculated as |1 - C_predicted_/C_observed_|, where C stands for correlation. The matrix representation of the normalized difference (Fig. 2A, third row) indicates that the model-based prediction was significantly more accurate in anesthesia than in wakefulness (Fig. 2B, left). Likewise, all pairwise Spearman correlation between observed and predicted correlation increased from 0% desflurane to 2, 4, and 6% desflurane (Fig. 2B, right) confirming that the model predicts correlation better in anesthesia than in wakefulness.

**Figure 2.**
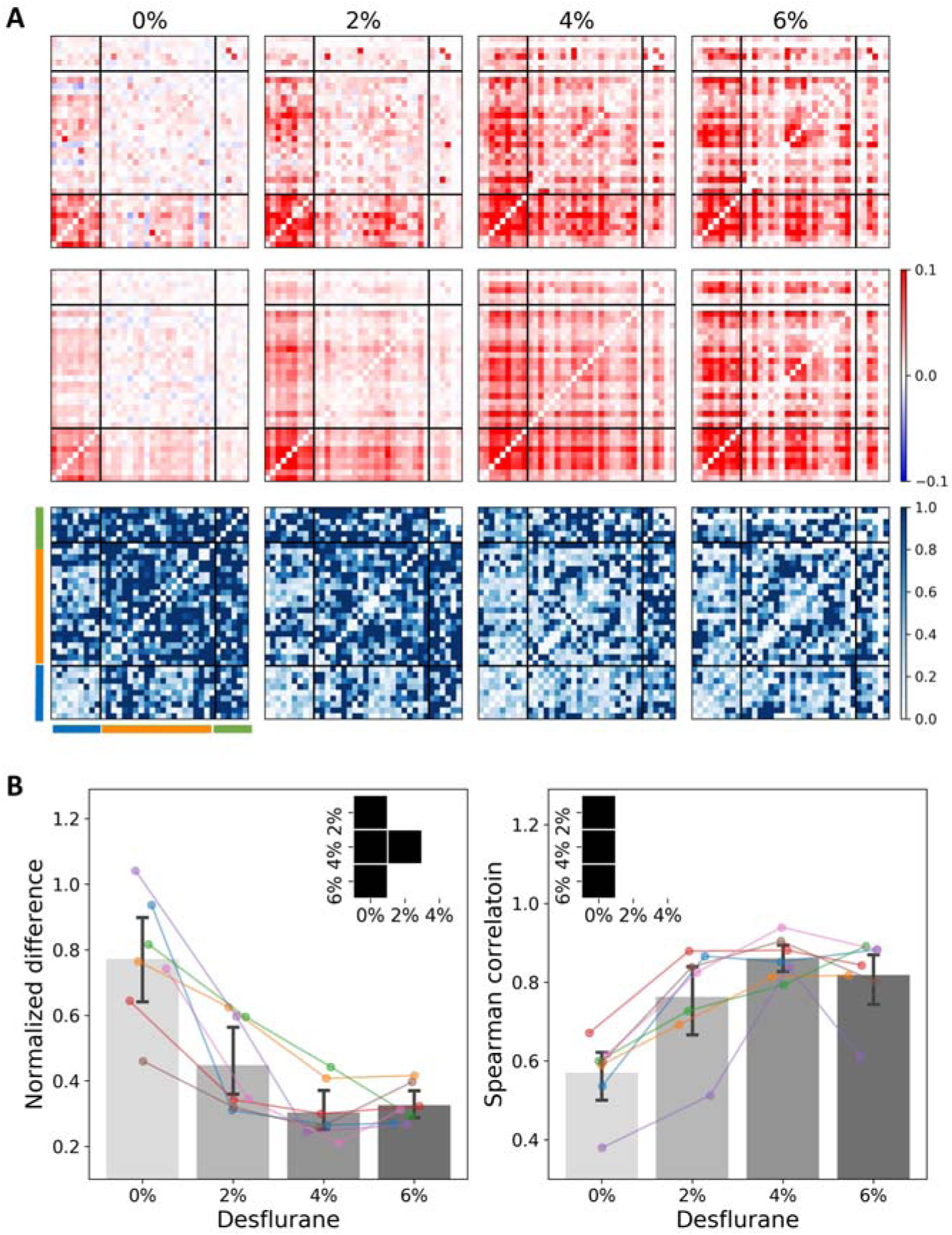
Pairwise correlation of spontaneous activity in four conditions. (A) First row: pairwise correlation estimated from experimental data of a representative animal. Second row: pairwise correlation predicted from a spike raster model that matches measured values of population coupling, firing rate, and population firing rate distribution in the same animal. Third row: normalized difference between observed and predicted correlation. Horizontal and vertical black lines divide neurons into three groups (blue: HPHF, orange: LPHF, green: LPLF); abbreviations are defined in Fig.1. Color bar scale indicates correlation value (first and second rows) and normalized difference (third row). (B) Mean normalized difference (left) and mean Spearman correlation (right) show better model prediction of correlation in anesthesia than in wakefulness. The seven color lines indicate data from different animals. Error bars indicate CI 95% across animals. The insets show pairwise statistically significant difference among conditions at p < 0.05. In all panels, 0%, 2%, 4% and 6% indicate inhaled concentration of desflurane.

### 3.4 Altered magnitude and timing of visual response

We explored the FR response to visual flash in the three neuronal groups under anesthesia and wakefulness. Consistent with the results of spontaneous activity, flash-induced FR was highest in HPHF and lowest in LPLF neurons at 0% desflurane (Fig. 3A, first row). In HPHF and LPHF neurons, the early increase in FR (20-200ms) was followed by a distinct suppression (a dip right after the early response) and late response activity (> 200ms). HPHF neurons showed a particularly prominent peak in the early response near 100ms. The late response persisted the longest in LPHF neurons; the time points where statistical significant response terminated were 190, 440, and 130ms, for HPHF, LPHF, and LPLF neurons, respectively. In addition, an oscillatory pattern was clearly seen in LPHF neurons.

**Figure 3.**
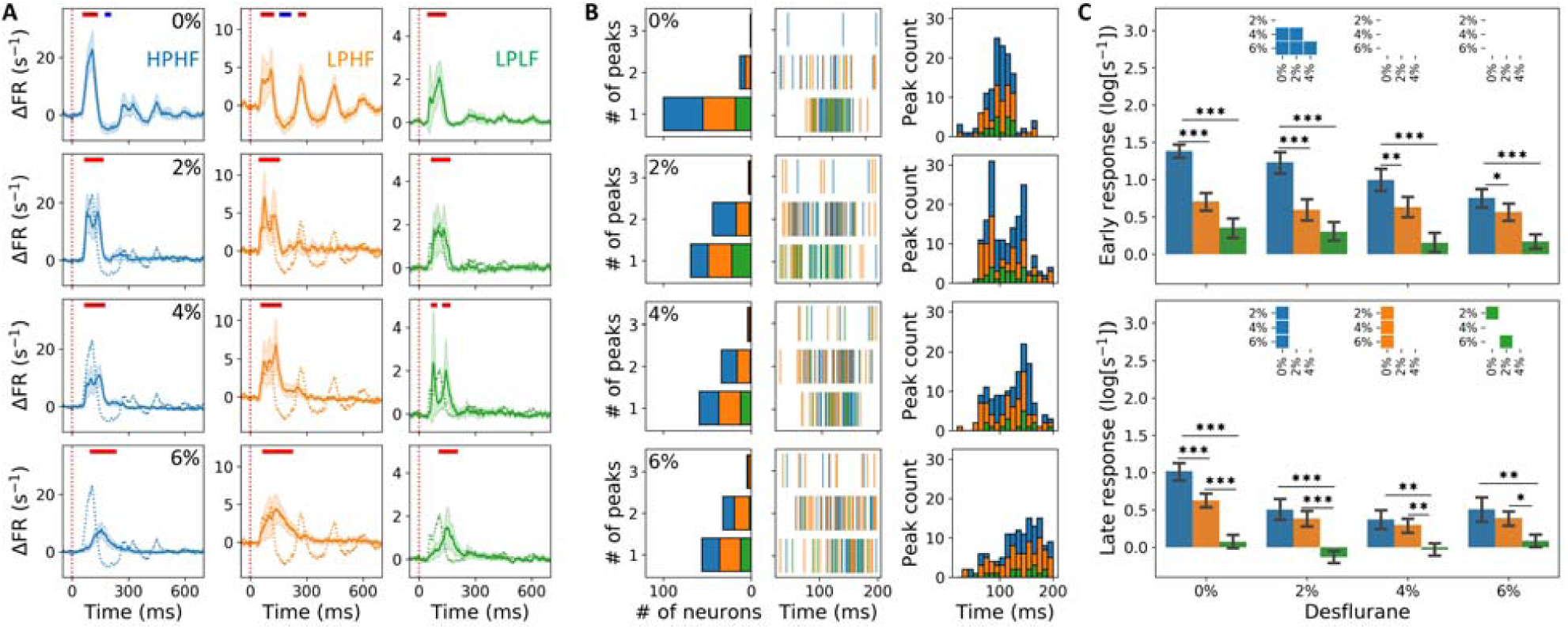
Neuronal response to visual flash stimulation in different anesthetic conditions. (**A**) Trace of spiking activity averaged across neurons: First column: HPHF (blue), second column: LPHF (orange), and third column: LPLF (Green). Flash is delivered at 0ms (vertical red dotted line). Relative spike activity compared to prestimulus baseline (FR minus the mean of FR in -1000 to 0ms) is represented. Shaded area represents CI 95% from the respective neurons. The horizontal color bars in each panel indicate significant difference (p < 0.001) of FR at time point compared to prestimulus baseline (red: increase, blue: decrease). (**B**) Change of spike timing under anesthesia. The number of spike rate peaks within 20-200ms were counted for each neuron and pooled in stacked bar graphs for three neuron groups (first column). The same color code for neuronal groups as in (A) is used. The time of peak of individual neurons is displayed (second column). Stacked histogram of the peak times indicate a significant change of spike timing by the anesthetic (third column). (**C**) Relative firing activity at the peak of the early (20-150ms, above) and late (200-400ms, below) response. Error bars indicate CI 95% across neurons within each group. Insets indicate statistically significant difference between pairs of anesthesia levels. Pairwise comparison between the three groups was also made at each level of anesthesia.

In anesthesia, both the suppression and late response were diminished (Fig. 3A, rows two to four), whereas the early response remained significant in all anesthetic conditions consistent with our former study [12]. However, the spike timing of HPHF and LPHF neurons within the early response (20-200ms) was altered such that the single FR peak in wakefulness was split into two peaks at 2% desflurane. This is further illustrated after smoothing each spike train with 20ms moving average (Fig. 3B). The two-peak pattern was most frequently seen in the HPHF group (61.4% of two-peaks were in HPHF neurons), which is large considering that these neurons were the least numerous (55 out of 251). From 2% to 6%, the two-peaks pattern diminished and spike timing was delayed (Fig. 3B, rows three and four).

We also quantified the magnitude (FR at peak response) of the early and late response (20-150ms and 200-400ms, respectively) and examined their changes across different levels of anesthesia (Fig. 3C). Only HPHF neurons showed a statistically significant decrease in early response from wakefulness to anesthesia (upper panel in Fig. 3C). The late response was clearly attenuated HPHF and LPHF neurons.

In summary, the latter findings confirm that desflurane anesthesia generally suppresses the late response of neurons more than their early response. In addition, the magnitude and timing of the early response of HPHF neurons are substantially altered.

## Discussion

The primary goal of this study was to test if anesthesia differentially affects neurons depending on their intrinsic firing properties. Firing rate and population coupling are thought to be important, invariant properties of neurons [1,2]. It has been reported that firing rate and population coupling of neurons are positively correlated [4], but we found that in wakefulness, many of high-firing neurons also disconnected from population. Therefore, different from previous studies in which neurons were classified by either firing rate or population coupling alone, we classified neurons into three groups based on both firing rate and population coupling, and investigated how anesthetic affects spike activity of these neuron groups.

As anesthesia deepened, the population coupling and firing rate of each neuron became increasingly correlated, meaning that most high-firing neurons acted as choristers and low-firing neurons acted as soloists. Moreover, the population coupling of all neurons significantly increased in anesthesia. The spikes of individual neurons are constrained to population activity; they only fire when the population fires, reducing the chance for processing a diverse content of sensory information; in other words, information capacity of the neural network becomes very low [13]. Considering the fact that soloists respond to sensory stimuli in a selective and precise manner [4,6], one could surmise that the absence of the actively firing soloists in anesthesia could be responsible for the disruption of conscious sensory information processing. This proposition is consistent with anesthetic-related changes in large-scale brain activity such as electroencephalogram, which exhibits slow oscillation, low-frequency synchronization (0.1-5 Hz), or suppressed signal complexity as typical signatures of anesthetic-induced unconsciousness [14–16].

Using a spike raster model, we also estimated the extent to which pairwise correlation of neurons’ firing could be predicted by their population coupling and firing rate distribution. We found that the model prediction was more accurate in anesthesia than in wakefulness. This means that during anesthesia there was little information in the neuron-to-neuron correlation in excess of that of which the population had. In the wakeful state, the prediction of pairwise correlation was less precise and the rank correlation between observed and predicted pairwise correlations was low suggesting a greater diversity of neural communication independent of population activity.

At critical levels of anesthesia, a disconnection between firing rate and population coupling was also observed. Namely, firing rate but not population coupling changed between 4 and 6% desflurane. Based on behavioral observation, i.e. the righting reflex, animals appeared conscious at 0 and 2% desflurane and unconscious at 4 and 6% desflurane. This suggests that population coupling indexed the anesthetic state of consciousness better than firing rate.

Chorister neurons have been reported as responsive [1] but non-selective to the sensory stimuli [4,6]. In our study, choristers showed the strongest response to visual stimulation among the three neuron groups. This may be related to the choristers’ stronger functional interconnection, as we indeed found. Likewise, Okun et al., observed that choristers received more synaptic inputs from their neighbors, suggesting that chorister-soloist characteristics could be attributed to anatomical connectivity structure and therefore, were invariant properties [1]. Consistently, we found that the rank of population coupling was conserved across the anesthetic conditions including wakefulness both in resting and stimulation-induced activities. Thus, our observations extend the generality of Okun’s finding suggesting population coupling is invariant with respect to different behavioral conditions and determined by structural connectivity.

Anesthesia disrupts sensory processing as a component of causing unconsciousness. However, studies using stimulation-evoked potentials have reported that primary sensory cortex remains responsive to sensory stimuli even under moderate level of anesthesia [12,17,18]. It has been postulated that conscious experience is more correlated with long-latency, sustained responses of high-order cortical areas, which integrate neural activity beyond the primary sensory regions [19]. The long-latency evoked potentials and neuron firings are thought to reflect reentrant processing, which appear to be preferentially suppressed in association with anesthetic-induced unconsciousness [12,18]. The spike response in our study gave similar results; the late response of high-firing neurons was significantly lower in the anesthetized than in the wakeful state. Moreover, in wakefulness, high-firing soloist neurons exhibited longer latency spike activity (∼440ms) than other groups and uniquely exhibited a distinct slow oscillatory pattern. In contrast, high-firing chorister neurons showed a large early response and early attenuation of their late response.

At variance with our previous study [12], the early flash-induced spike response was not always preserved, as the response magnitude of high-firing chorister neurons progressively decreased in deeper anesthesia. The early response of these neurons also split into two peaks at the lowest anesthesia level. This suggests that classifying neurons into functional groups based on their firing rate and population coupling can reveal additional details of the state-dependent alterations in the neurons’ sensory response that are not discernible when pooling neurons into a single population.

In summary, anesthesia promoted a strong association between firing rate and population coupling, suppressing population-independent spike activity. Classification of neurons revealed that the effect of anesthesia on stimulation-induced activity is distinct between neuronal populations. This study advanced the neuron-level understanding of how anesthesia affects the primary sensory cortex and how it disrupts sensory processing.

## Acknowledgements

Research reported in this publication was supported by the National Institute of General Medical Sciences of the National Institutes of Health under award number R01-GM056398 and the Center for Consciousness Science, Department of Anesthesiology, University of Michigan Medical School, Ann Arbor, Michigan, USA. The content is solely the responsibility of the authors and does not necessarily represent the official views of the National Institutes of Health. The authors thank members of the Center for Consciousness Science, University of Michigan Medical School, for valuable comments and Kathy Zelenock, MS for her assistance in laboratory operations and manuscript editing.

